# Adult Marine Annelid *Platynereis dumerilii* Chemically Stunt the Growth of Juveniles

**DOI:** 10.64898/2026.04.30.721953

**Authors:** Victoria C. Moris, Paula Schirrmacher, Sophie Potter, Martha Tickle, Ryan Squire, Jörg D. Hardege

## Abstract

Within species, individuals of the same age can differ in size. Previously, parental genetics, nutrition, space, and social interactions have been suggested to explain different growth rates. However, direct effects of larger individuals on the physiology and growth of smaller individuals are poorly understood. In this study, we investigated how larger individuals of the marine worm *Platynereis dumerilii* can impact the growth of smaller conspecifics. Comparing growth distributions in communally and individually reared worms, we show that larger worms suppress the growth of smaller ones. Furthermore, we were able to demonstrate that this suppression is chemically mediated. The chemical cue does not originate from faeces but is water soluble, stable for several days and smaller than 3 kDa. Our findings highlight the importance of non-reproduction related chemical signalling, showing evidence that dominant individuals can chemically suppress the growth of their conspecifics. This study provides new insights into how hierarchy can be established and maintained in a population and is particularly relevant for the growing community studying this model species.

## Introduction

Why, even in similar conditions, siblings grow to different sizes, is a fundamental puzzle of life. Differences in growth rates can often be explained by parental genetics, nutrition and space (Sibly et al., 2005). Indeed, animals interact with others, inevitably competing for space, food and partners in dense, multigenerational communities. Competition and social interactions often trigger aggressive behaviours and dominance hierarchies (Briffa and Sneddon, 2007; Drews, 1993; Gherardi and Atema, 2005), which may also lead to growth variation. Several decades of research on the social aspects of population dynamics provide evidence of social dominance often mediating access to mating and food (Briffa et al., 2015; Tibbetts et al., 2022). Thereby, hierarchical systems (or dominance) can be beneficial for both dominant and subdominant individuals, asserting their status whilst minimising costly fights (Alcala et al., 2019; Tibbetts et al., 2022). However, direct effects of dominant individuals on the physiology and growth of subdominants are poorly understood. Population density is known to affect metabolic rates (DeLong et al., 2014). Furthermore, it has recently been suggested that conspecific chemical cues, not food competition, could drive metabolic suppression in bryozoans (Lovass et al., 2020).

The ragworm *Platynereis dumerilii* (Nereididae), is a promising group to investigate hierarchy systems and impact of dominant individuals on the growth of subdominants. Populations comprising individuals of widely different sizes have been observed in laboratory cultures of *P. dumerilii* (Kuehn et al., 2019; Kuehn et al., 2022). *P. dumerilii* as a model species, has been the focus of a growing scientific community (Özpolat et al., 2021), particularly studying neuro-biological and evolutionary research (Jékely et al., 2008; Williams and Jékely, 2019), as well as for ecology and toxicology (Hardege, 1999; Hutchinson et al., 1995). In the wild, *P. dumerilii* play a fundamental role in marine and estuarine food webs (Lawrence and Soame, 2009). In the wild as well as the laboratory, immature *P. dumerilii* dwell in mucous tubes, interacting with conspecifics to forage (Özpolat et al., 2021). Its segmented body and regular growth (up to one segment per day, throughout juvenile stages (Hauenschild and Fischer, 1969)) provides a straightforward method for quantifying size/growth as a proxy for dominance.

Growth, maturation and reproduction in Nereididae are thought to be regulated by at least two hormonal systems: sesquiterpenoid methylfarnesoate, a juvenile hormone called Nereidine (Schenk et al., 2016), and a gonadotrophic maturation activity. Nereidine promotes growth and regeneration of lost segments in young animals (Golding, 1967). While inhibiting sexual maturation, it also leads to the accumulation of oogonia into the coelom (Franke and Pfannenstiel, 1984; Golding, 1967). In contrast, a gonadotrophic maturation activity, proposed to originate from mature female ganglia, has been suggested to stimulate the growth of oocytes (Lawrence and Soame, 2009). The relative levels of juvenile and maturation related hormonal activity are likely balanced across life stages (see scheme Fig. 1).

**Fig. 1.**
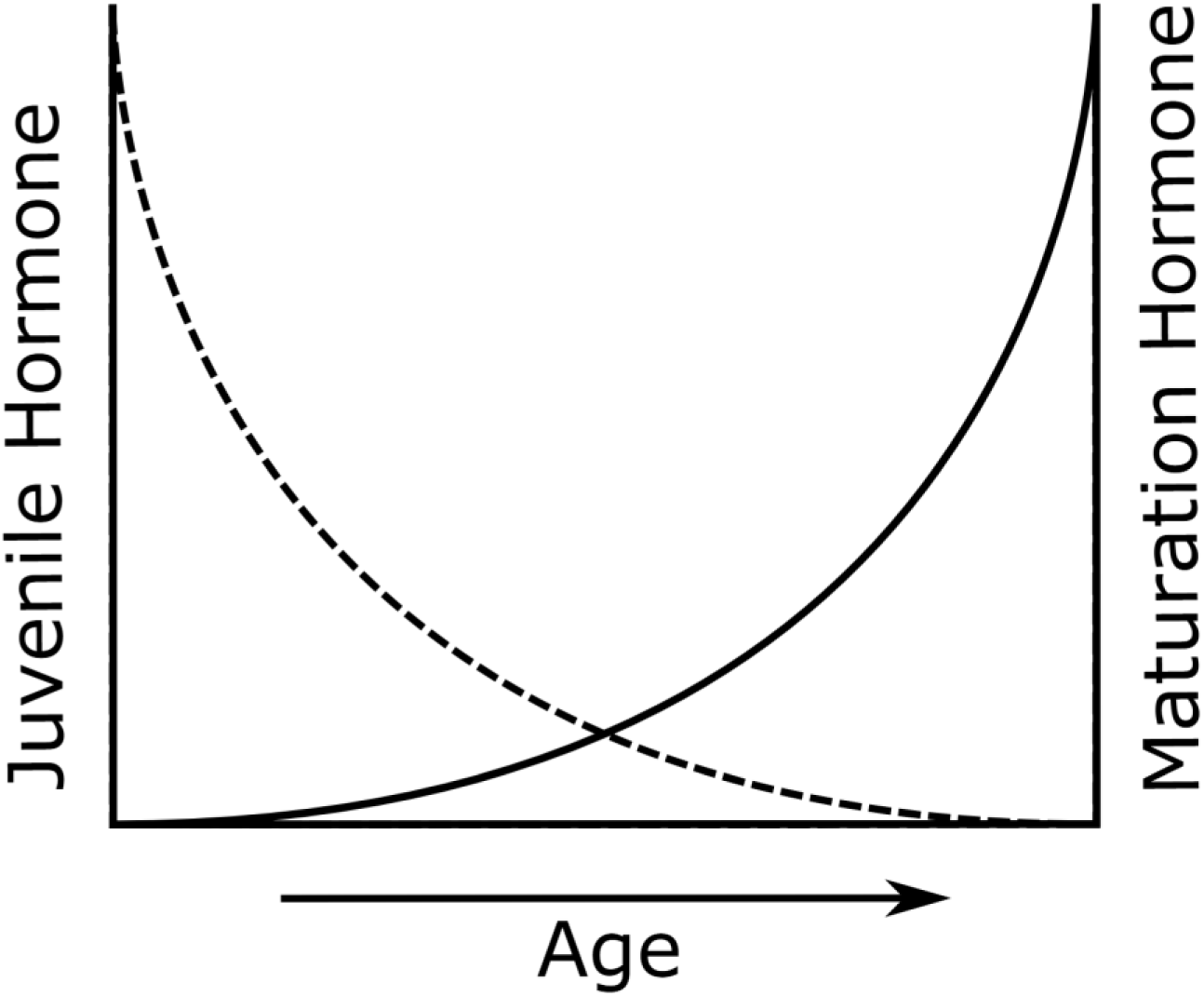
Scheme of the proposed balance of juvenile (dashed line) and maturation hormone (solid line) in Nereidids.

In this study, we investigated mechanisms underlying hierarchy systems in in the annelid worm *Platynereis dumerilii*. First, we examined growth distributions in rearing conditions with and without interactions among conspecifics. Separating large and small worms physically, whilst allowing for water exchange between the groups, we then evaluated the effect of chemical exudates from large worms on the growth of smaller ones. We report a significant growth reduction triggered by the exposure to water from larger worms. Finally, we showed that the growth suppression is mediated chemically and determined some chemical characteristics of the signaling cue by assessing its stability over time and investigating the approximate size of the compound. This study intends to shed light on non-reproduction related chemical communication in annelids and improves our understanding of the general mechanisms underlying social hierarchies.

## Materials and Methods

We investigated the growth suppression in *P. dumerilii* with a set of three experiments (summarized in Fig. 2). First (Fig. 2a), we compared the growth of small worms in individual and communal rearing conditions with comparable access to food and space. Next (Fig. 2b), we studied the growth of small worms exposed to chemical (and visual) cues from large worms, excluding direct interactions by separating the groups with fine gauze (dashed line in Fig. 2b). Finally (Fig. 2c), we carried out experiments with small and large worms in separate tanks, exposing juveniles to faeces from adult worms, water from adult worms and size fractions of adult worm water (<3 kDa and>3 kDa). All materials and methods are covered in more detail below.

**Table 1.** Number of worms per tank and number of tanks in the three experiments at T1 (0 weeks) and T2 (2–3 weeks of treatment).

| Experiment | Condition | # Weeks | Worms per tank | # Tanks | N T<sub>1</sub> | N T<sub>2</sub> |
| --- | --- | --- | --- | --- | --- | --- |
| a. Indiv vs. Communal | Individual | 6 | 1 | 50 | 50 | 50 |
|  | Communal | 6 | 25 | 2 | 50 | 50 |
| b. Lab | Small vs large (adults) | 3 | 25 | 7 | 175 | 161 |
|  | Small vs small (control) | 3 | 25 | 6 | 150 | 137 |
| b. F1 field-derived | Small vs small | 2 | 25 | 3 | 75 | 29 |
|  | Small vs medium | 2 | 25 | 3 | 75 | 34 |
|  | Small vs large | 2 | 25 | 3 | 75 | 52 |
|  | Medium vs small | 2 | 25 | 4 | 75 | 59 |
|  | Medium vs medium | 2 | 25 | 3 | 75 | 67 |
|  | Medium vs large | 2 | 25 | 3 | 75 | 65 |
| c. Faeces | Adult faeces | 3 | 25 | 3 | 75 | 70 |
|  | Water (control) | 3 | 25 | 3 | 75 | 70 |
| c. Water | Adult water | 3 | 50 | 9 | 450 | 194 |
|  | Water (control) | 3 | 50 | 3 | 150 | 78 |
| c. Size | Size < 3 kD | 2 | 20 | 2 | 40 | 31 |
|  | Size > 3 kD | 2 | 20 | 2 | 40 | 26 |
|  | Water (control) | 2 | 20 | 2 | 40 | 32 |

**Fig. 2.**
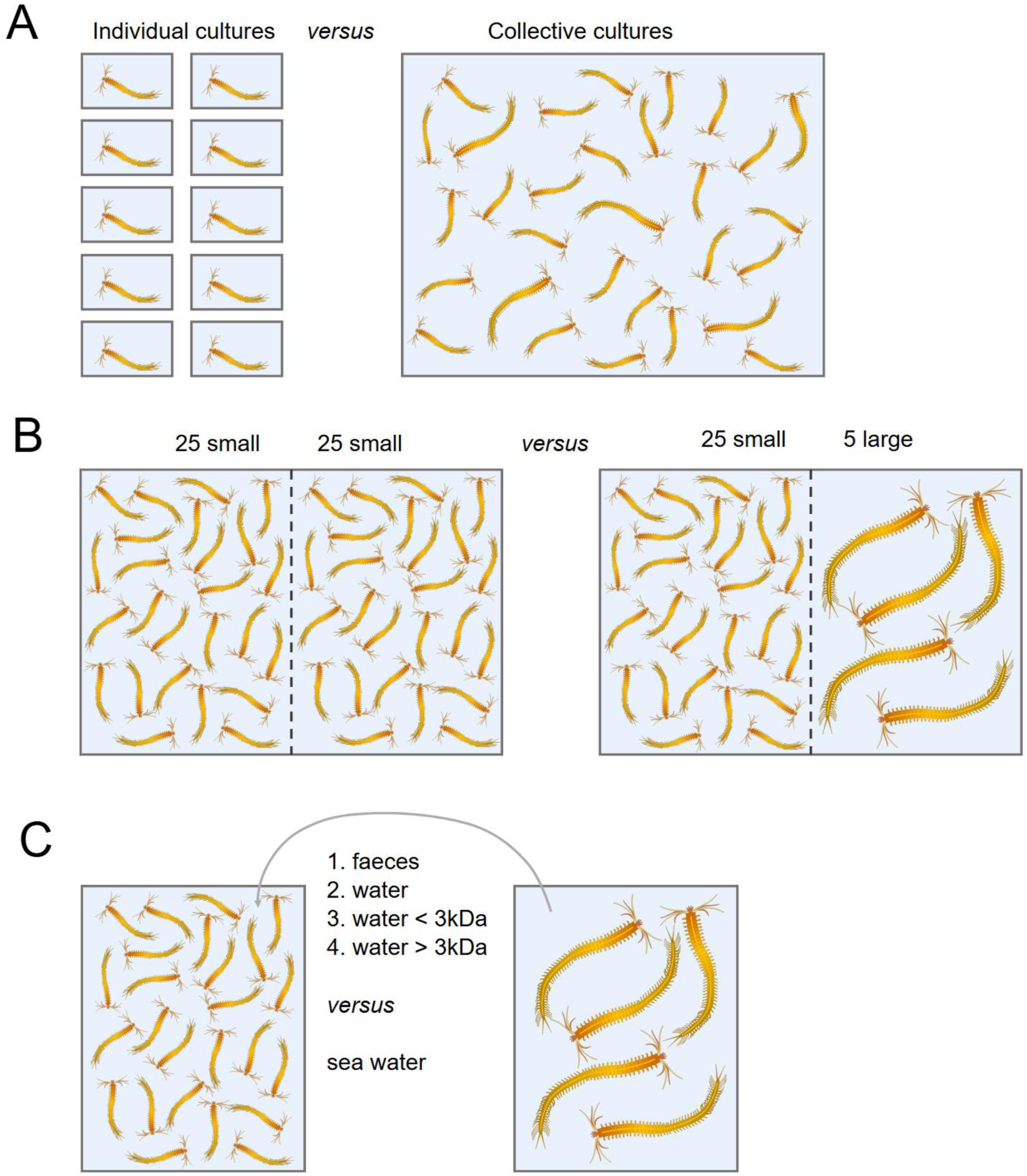
Scheme of the tank set-ups to explore growth suppression in *P. dumerilii*. In the first set of experiments (A), we compared the growth of worms in individual and communal rearing conditions. Next (B), we studied the growth of small worms exposed to chemical cues from large worms, using small worms as control. The two groups of worms are separated by fine gauze (dashed line) to exclude direct interactions. Finally (C), we examined the growth of juveniles exposed to adult *P. dumerilii* faeces, water and water size fractions <3 kDa and>3kDa, compared to sea water as control, from separate tanks (excluding potential visual stimuli of large worms).

### Worm Collection and Husbandry

This study uses the F1 offspring from field-collected *Platynereis dumerilii* individuals (“F1 field-derived”) maintained in the laboratory for a single generation, as well as an established laboratory culture (kindly donated by Prof. G. Jékely). The field collection of polychaete worms *Platynereis dumerilii* was carried out at Hinkley Point, near Bristol, on 4th November 2009. Molecular phylogenetic analyses by (Calosi et al., 2013) confirmed the classification of the *Platynereis* species at this location. At the University of Hull, both established laboratory and the F1 field-derived cultures were kept at 18^∘^ C in 20 cm × 20 cm tanks containing 1.6 L filtered natural sea water with an average salinity of 35 ± 0.5. The tanks were kept aerated and in a light/dark cycle of 16 h/8 h. To mimic the lunar cycle, a low light lamp was introduced during full moon, similar to the culture set up described by (Kuehn et al., 2019). Worms were fed twice a week in excess. *Tetraselmis marina* phytoplankton culture was the major food source for small worms, larger juveniles and adult worms were also fed defrosted spinach. Excess food and faecal material were removed twice a week using a pipette. The tank water was changed once per week.

### Individual vs. Communal Rearing (a)

We compared the segment growth of small worms reared individually and small worms reared in communal conditions using worms from the established laboratory culture (Fig. 2a). As both communally and individually reared worms originated from the same clutch, genetic differences are not expected to influence the outcome. All specimens used in the experiment were derived from a single clutch initially reared communally. For individual rearing, small worms (16-24 segments, 2 sets of 25, N=50, 50 small tanks, ([Table1]) were placed in separate cylindrical containers with 17 mm radius, containing approx. 36 mL of filtered natural sea water and 0.5 mL *Tetraselmis marina*. For communal rearing, small worms (16-34 segments, N=50, 1 tank divided into halves) were placed in a tank containing 1.6 L natural sea water. The tank (28 cm × 16 cm) was divided by fine gauze into two equal parts with 25 worms on either side (see set-up Fig. 2b) to ensure an equal distribution of worms and reduce fighting. The cultures were fed ad libitum and cleaned as described above (worm collection and husbandry section), except that the water was only changed every 2 weeks to enable build-up of chemical cues and metabolites excreted from worms. Over the course of 6 weeks, their size (number of segments) was measured in 2 week intervals.

### Size Dependent Growth Suppression (b)

Since we observed different average growth in worms reared individually and communally, we next addressed the question whether the observed growth suppression in communal rearing conditions is due to visual/ social interactions or whether it is chemically induced. This experiment was carried out with worms coming from the established laboratory culture. The worms were placed in 28 cm × 16 cm tanks containing 1.6 L sterile filtered water (0.2 μm filter size). The tank set-up (as visualized in Fig. 2b) was adapted from Cobb and Tamm (1974): The tank was separated into equal parts by a fine gauze (250 μm). This ensured a constant exposure to freshly released chemical cues from worms of the opposite tank side whilst excluding social interactions between worms from opposite sides. The segment growth of groups of 25 small worms (16-24 segments) was compared between exposure to water from 5 large worms (≥ 80 segments, ‘adult worms’) and exposure to other small worms as control (Fig. 2b). At 80 segments, worms mostly stop growing new segments and start growing the size of their segments (Fischer and Dorresteijn, 2004). Adult worms were kept in groups of 5 due to lack of space. The exposure to worms of the same size on the other side of the gauze served as control experiments. The cultures were fed ad libitum and cleaned as described above (worm collection and husbandry section), but the water was only topped up, not changed, to allow for the build-up of chemical cues. The growth of the worms was recorded after 3 weeks. Seven tanks of small worms (n=25 in each) were exposed to adult worm water, 6 tanks, used as control, contained small worms on both sides of the gauze (n=25 on each side of the gauze, 50 worms per tank, ([Table1])). For the analysis of small worms exposed to other small ones, we considered only the worms from one side, as we did when exposed to large worms.

We repeated this experiment with the F1+ field-related culture to confirm the results we observed with the established laboratory culture. Building on the protocol with laboratory culture worms, we exposed 25 small worms to a) 25 other small worms (control), b) to 25 medium worms, c) to 5 large worms. We also exposed 25 mediums worms to a) 25 small worms, b) 25 medium worms, and to c) 5 large worms. Each condition was performed with 3 tanks ([Table1]). Hereby, small worms were defined as 16 to 40 segments long, medium-sized worms were 40 to 65 segments and large worms were longer than 80 segments. Due to working with specimens originating from field collection, the culture was smaller, making it more difficult to use worms with the same number of segments for the start of experiments. The cultures were fed ad libitum and cleaned as described above (worm collection and husbandry section). Before and after the 2 week experimental period, the size of worms (segment length) was recorded.

### Chemical Characteristics of Growth Suppression (c)

To determine chemical characteristics of the growth suppression cue, we carried out three experiments (Fig. 2c): we tested whether the cue originated from faecal material of adult worms or was water-borne, and then explored the effect of size fractions of water from adult worm tanks.

To explore the effects of faecal material alone, we used the same gauze-separated tank set-up as above, 25 worms were placed on one side of the gauze. The other side of the tank contained faecal pellets or was left empty as control. Experiments were run in triplicate (3 tanks per condition, containing each 25 worms, ([Table1])). Faecal pellets were collected from 5 large worms (housed in a separate tank) every three days, rinsed with artificial seawater and placed into the experimental tanks. The compact nature of *P. dumerilii* faeces (Gambi et al., 2000) allowed us to handle faecal material with a pipette without it losing its integrity. Pellets were rinsed by adding them to artificial seawater in a 50 ml Eppendorf tube and decanting the water three times.

Having addressed whether the chemical growth suppression cue originates from faeces, we next explored whether a component of the adult worm water could be responsible for the observed effect. We hence exposed 50 small worms (16-24 segments) to filtered adult worm water (0.2 μm, to exclude faecal material) or artificial seawater as control. Experiments were carried out in 28 cm × 16 cm tanks with 1.6 L water of the respective condition. Worms received their usual diet in excess twice a week. The water was changed every 3 days. The growth of the worms was recorded after 3 weeks. Nine tanks received the adult worm water, whilst three tanks, used as control, received clean culture water; each tank contained 50 worms ([Table 1]).

To determine chemical characteristics of the growth suppression cue, 20 small worms (20-48 segments) were exposed to filtered water (0.2 μm, to exclude faecal material) from adult worm tanks, which was fractionated with a 3 kDa filter (Milipore Amicon ultrafiltration membranes YM3 regenerated cellulose filters). A stirred ultrafiltration cell (Model 8003) was pressurised with nitrogen for the filtration. This resulted in two size fractions (<3 kDa and>3 kDa) of the molecules in the adult worm water, which was subsequently added to the treatment groups. The tank water of the small worms was changed every 3 days, whereby one group received water from the size-fraction smaller than 3 kDa and another group received water fractionated to contain only molecules larger than 3 kDa. The control group received clean culture water. The worms were fed their usual diet ad libitum (see worm culture and husbandry section above). Each condition was performed in duplicate (2 tanks containing each 20 worms, ([Table 1])). The segment growth was observed after 2 weeks.

### Statistics

Statistics were performed in R (R Core Team, 2019, version 4.3.3).

For experiment a, we analyzed whether growth trajectories differed between treatments using an analysis of covariance (ANCOVA) testing the homogeneity of slopes between conditions. Because individuals in the communal treatment could not be individually identified, the tank was considered the experimental unit; therefore, mean segment size (number of segments) per tank per time point was used in the analysis. We fitted a linear model with size as the response variable and time, condition (individual vs. communal).

For experiments b and c, we first calculated the mean segment length in each tank before and after exposure to the different treatments to account for the tank effect. The difference in segment length was then divided by the duration of the experiment (in weeks) to obtain the growth rate (segments per week) for each tank. Each tank-level mean was based on 20–50 individual observations. We compared the mean growth rates of the control tanks with those of the treatment tanks (e.g. adults, medium, large, adult water, size > 3 Da, size < 3 Da). When comparing two groups, we used the Student’s t-test when sample sizes were equal and group variances did not differ significantly, and the Welch’s t-test when sample sizes or variances differed. When comparing three groups, we used a one-way ANOVA, followed by Tukey’s post-hoc tests for pairwise comparisons.

## Results

### Growth Distribution (a)

Rearing 50 worms individually versus communally (Fig. 2a), we observed major differences in their growth distribution over the course of 6 weeks. Firstly, there was a large difference in median growth. Within 6 weeks, individually reared worms grew on average 14 segments (18 segments to 32 segments median length, Fig. 3a, Fig. 5, Fig. 4), whilst the average segment length remained unchanged in communally reared worms (30 vs 30 segments median length, Fig. 3b, Fig. 4). All 2 week intervals, except for 4-6 weeks, led to significant growth in the individually reared condition (Kruskal-Wallis test with pair-wise Dunn test and Bonferroni adjustment, 𝜒^2^= 91.2, 𝑃<0.001), whilst there was no evidence for significant growth in any of the 2 week intervals in the communal rearing condition (Kruskal-Wallis test, 𝜒^2^ = 6.1, 𝑃=0.11). Indeed, growth rates differed strongly between the two conditions (time × condition interaction, F = 9.96, p = 0.002; Fig. 4). Individuals raised alone showed positive growth over time (slope = 2.12), whereas those raised communally exhibited no net growth (slope = –0.20).

**Fig. 3.**
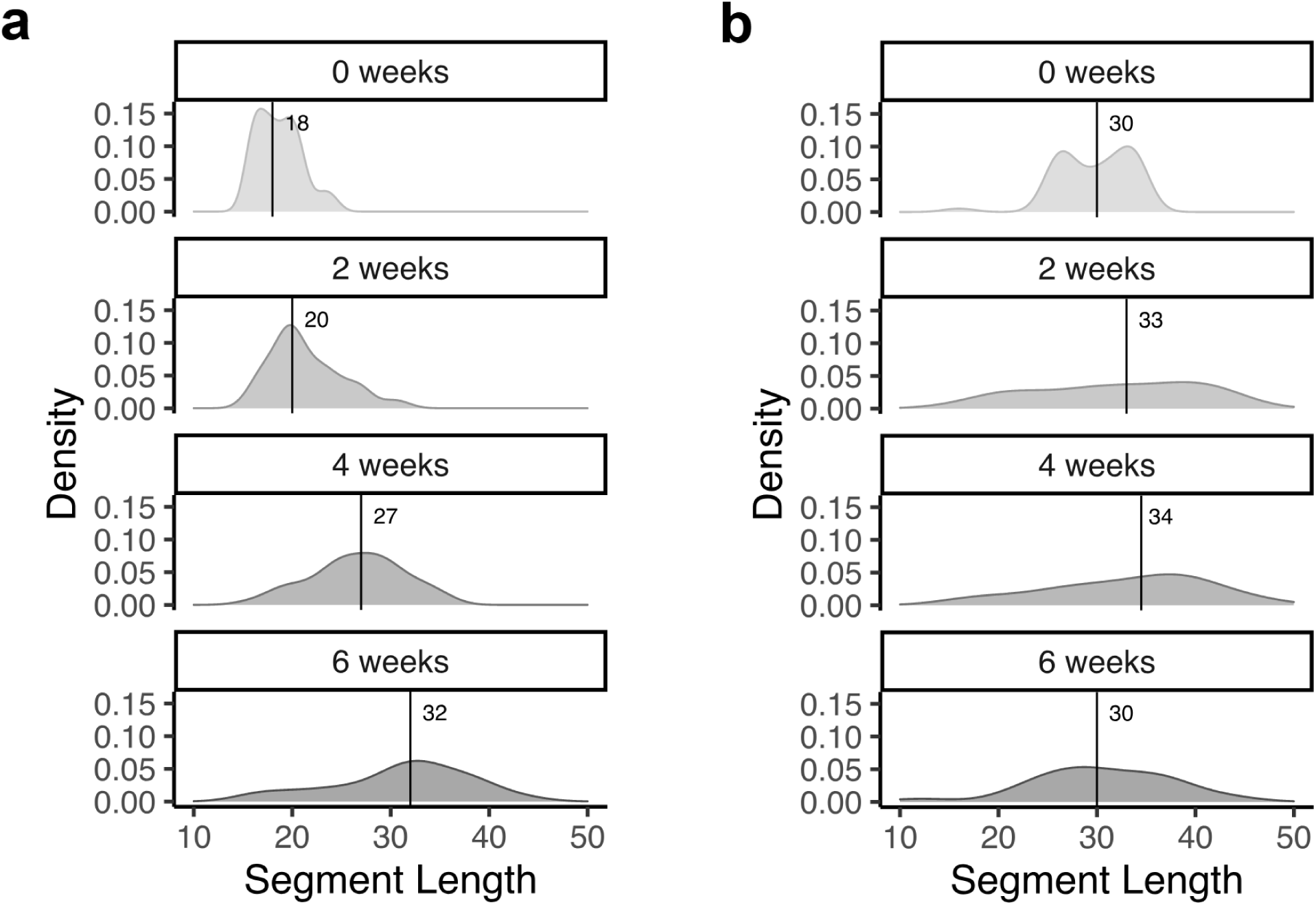
Density plots of the size distribution (in segment lengths) of individually (A) and communally (B) reared worms over the course of 6 weeks. The area under the density curve represents the probability of observing a specific segment length or range of segment lengths. Vertical lines and numbers display the median segment length at the respective timepoint.

**Fig. 4.**
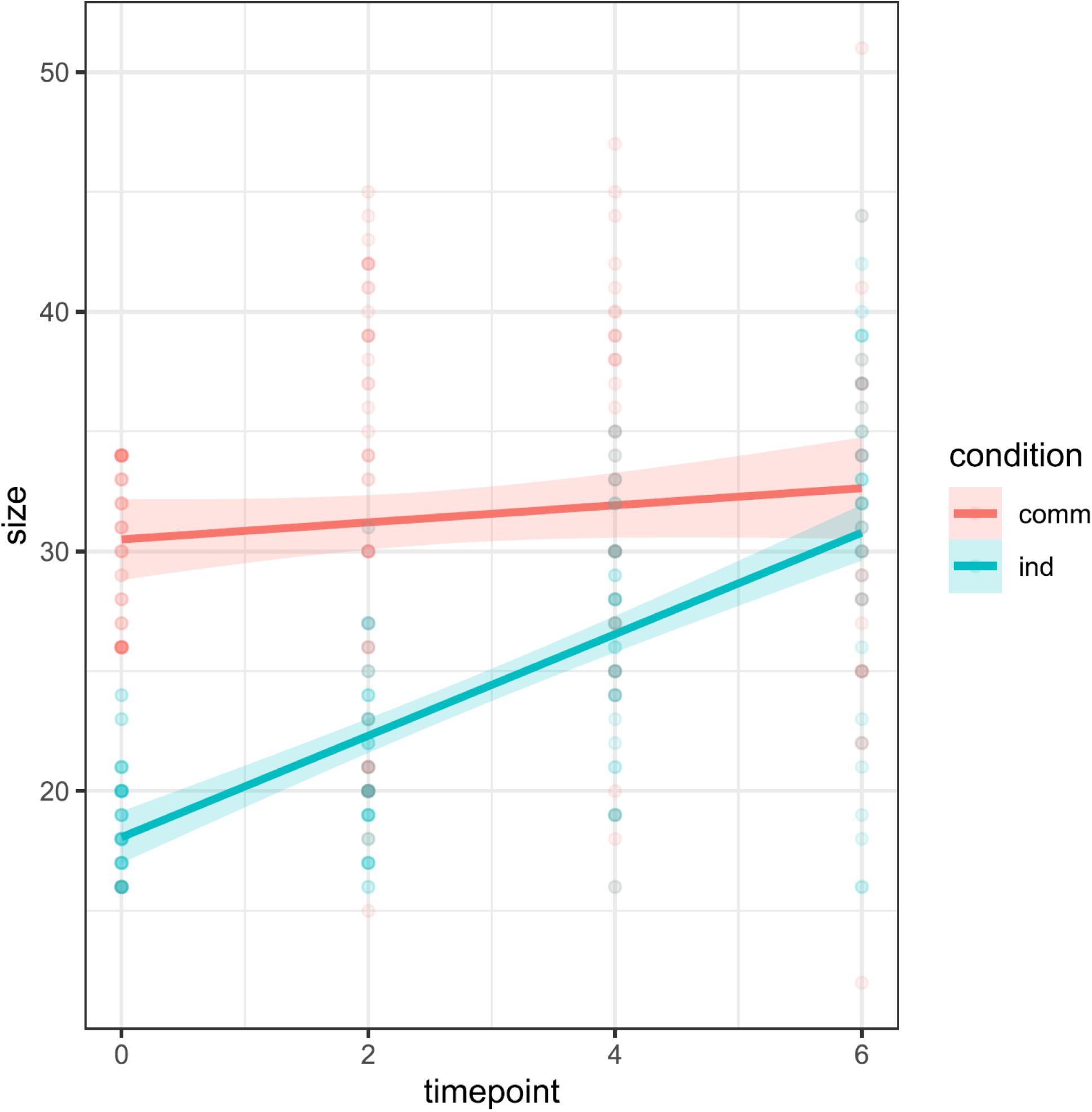
Growth trajectories for the two treatments (individual in green, and communal cultures in red). All individual measurements are plotted as semi-transparent points. Lines show the fitted linear models and shaded regions denote 95% confidence intervals. Because individuals in communal tanks could not be individually tracked, all measurements from communal tanks are shown; tank identity was accounted for in the statistical analyses.

**Fig. 5.**
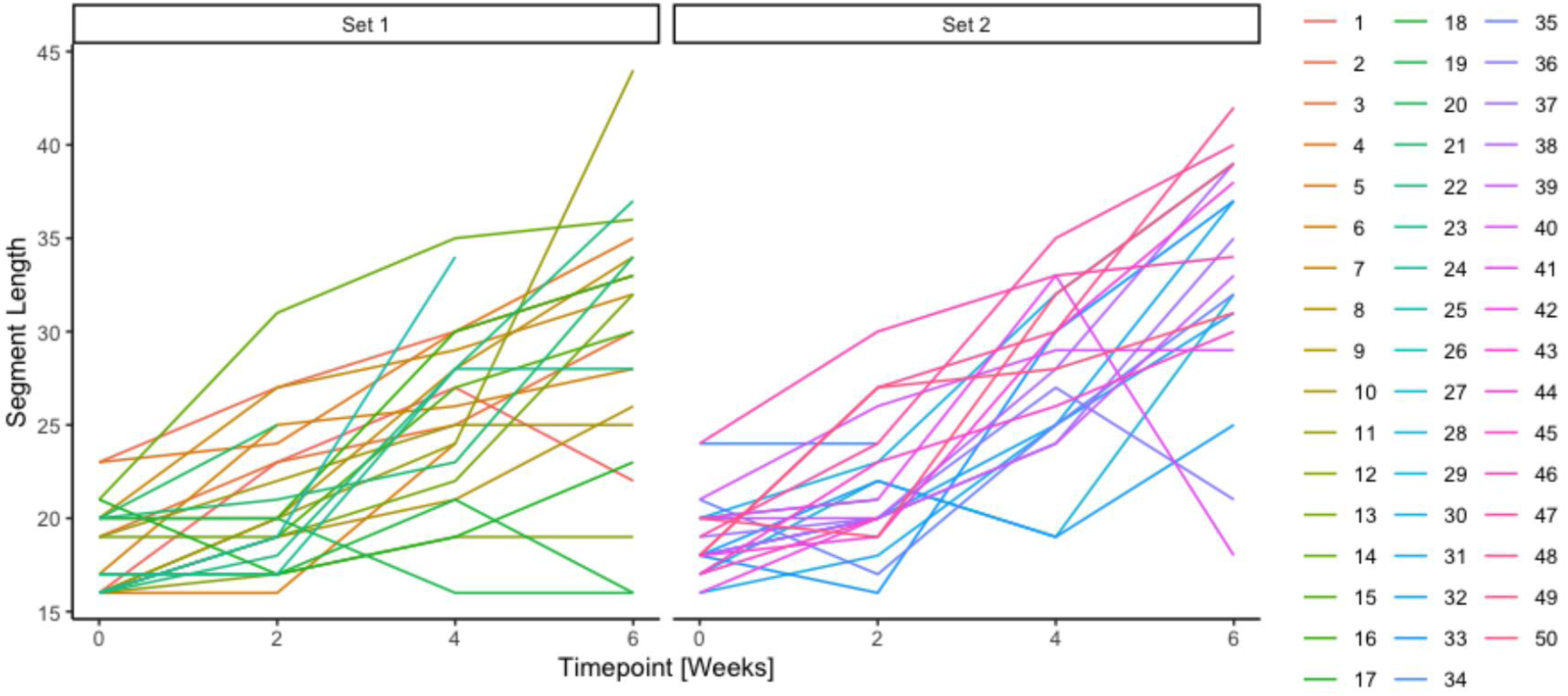
Segment lengths of *P. dumerilii* specimens reared individually over a period of six weeks. The samples were split into two sets for visual clarity; occasional drops in segment number are likely due to autotomy from handling.

Similarly for the two conditions, the density plots show clear changes in their distributions. Whilst the initial distributions are narrow and skewed (0 weeks), both final distributions are wider and normally distributed (Shapiro-Wilk normality test, 𝑊 = 0.961, 𝑃 = 0.174 for individual rearing; 𝑊 = 0.975, 𝑃 = 0.741 for communal rearing). However, we observed differences in the spread of sizes in individual versus communal cultures. At week 0, the size distribution showed multiple peaks in both conditions: three for individually reared specimens and two for communally reared ones (Fig. 3). These differences likely reflect the random selection of individuals for each experiment and illustrate the start of the experiments which originated from the same communal culture. The overall segment lengths were comparable between the two groups at the start (standard deviation 2.2 segments for individual and 3.7 segments for communal rearing condition). Over time, these distinct peaks in size distribution diminished in both conditions. After two weeks, individually reared worms displayed a single, narrow peak, indicating more uniform growth, whereas the distribution in the communal group broadened, with the initial peaks disappearing entirely showing a wide distribution across specimens (standard deviation 3.6 segments for individual, 8.3 for communal). The peak is less pronounced after four weeks of individual rearing. After 6 weeks, the difference in the spread of segment lengths evened out between the rearing conditions (standard deviation 7.0 segments in individual and 7.8 segments in communal rearing conditions).

Mortality was lower in the individual rearing conditions: 9 of the 50 worms died in individual cultures, whilst 23 of the 50 communally reared worms died/disappeared over the course of 6 weeks. Finally, some worms, particularly those in the communal rearing condition, showed a reduction in segment number over the course of six weeks, likely due to aggressive interactions or handling stress. Notably, we also observed a decrease in segment number in five worms reared individually (Fig. 5), where aggressive interactions were not possible. This suggests that the reductions were most likely caused by autotomy triggered by handling. However, since handling procedures were consistent across both individual and communal conditions, and the overall effect was minimal, we consider this impact to be negligible.

### Size-dependent Growth Suppression (b)

To determine whether the observed growth suppression in communal rearing depends on the size of the worms, we exposed groups of small worms to groups of large worms and other small worms (control), whilst physically separating the groups in a tank using fine gauze (Fig. 2b). This allowed us to focus on chemical cues, whilst excluding social interactions between the groups. Size categories were defined as follow: small worms were defined as 16 to 40 segments long, medium-sized worms were 40 to 65 segments and large worms were longer than 80 segments.

Using established lab cultured specimens, our results indicate that the growth rate (number of segments grown per week) of juveniles (small worms) exposed to adults (large worms) was significantly lower (mean growth rate= 2.5 +- 0.4 segments/week) than those exposed to same-size juveniles used here as control (mean growth rate= 3.8 +- 0.7 segments/week; Fig S1; Student t-test, df=11, t = 2.992, 𝑃=0.012). This suggests that large worms (adults) significantly stunt the growth of juveniles (Fig. 6; Fig S1).

**Fig. 6.**
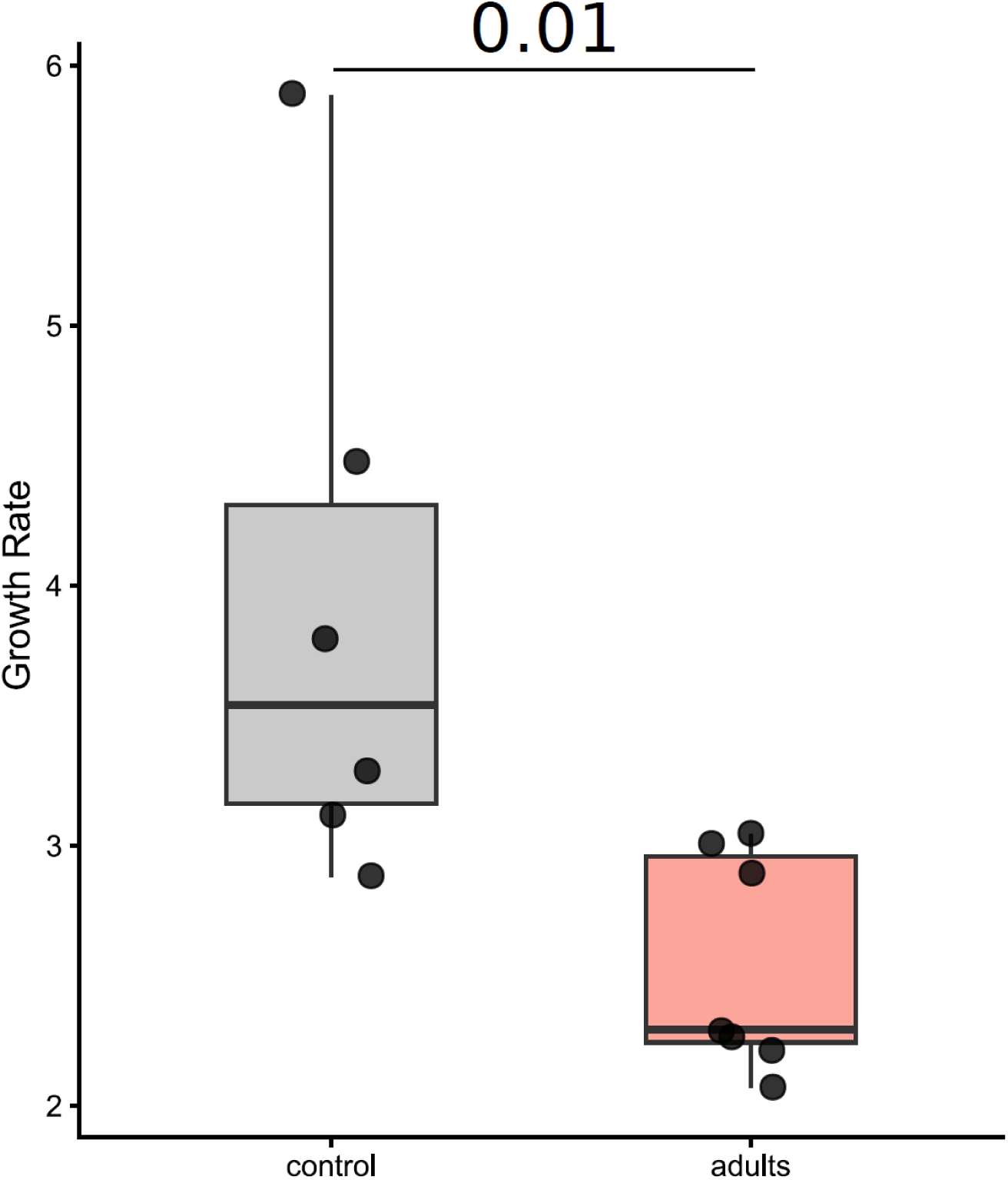
Growth rate of worms from established laboratory culture: small worms exposed to small worms (control) and to large worms (adults) over 3 weeks. The boxplots depict the median growth rate (segment length grown/week) with the first and third quartile of the distribution. Whiskers extend to 1.5 × the interquartile range; data beyond that range are defined as outliers and plotted individually. The p-value of the Student t-test comparing the two conditions is written between the two boxes.

We did not measure a significant effect of the condition (small, medium, large) to the growth of small worms (Anova, F=1.82, 𝑃= 0.24) although we observed that it seems slightly reduced when they were exposed to larger ones (mean growth rate= 0.26 +- 0.16 segments per week, Fig 7). This might be caused by the higher mortality of F1 field-related worms which caused the abortion of 2 of the 5 experiments per condition, leaving 3 repeats of each condition that were included in the analysis. In addition, the growth rates measured for these worms were lower than those of the lab culture. For example, small worms exposed to the same-size worms (control) had a growth rate of 0.67 +-0.44 segments/week (Fig 7; Fig S2) instead of 3.9 segments/ week that we measured in small worms from the lab culture. While we only had access to 35-segment worms when testing the adult water condition to small worms (Fig S2), we expected these worms to grow linearly, as observed for other small worms (Fig 3a and Fig 5). The fact that we observed a tendency to smaller growth for these 35-segment worms exposed to adult-conditioned water supports our previous observation that smaller worms (15–20 segments) exposed to adult worms also show inhibited growth (Fig. 6).

**Fig. 7.**
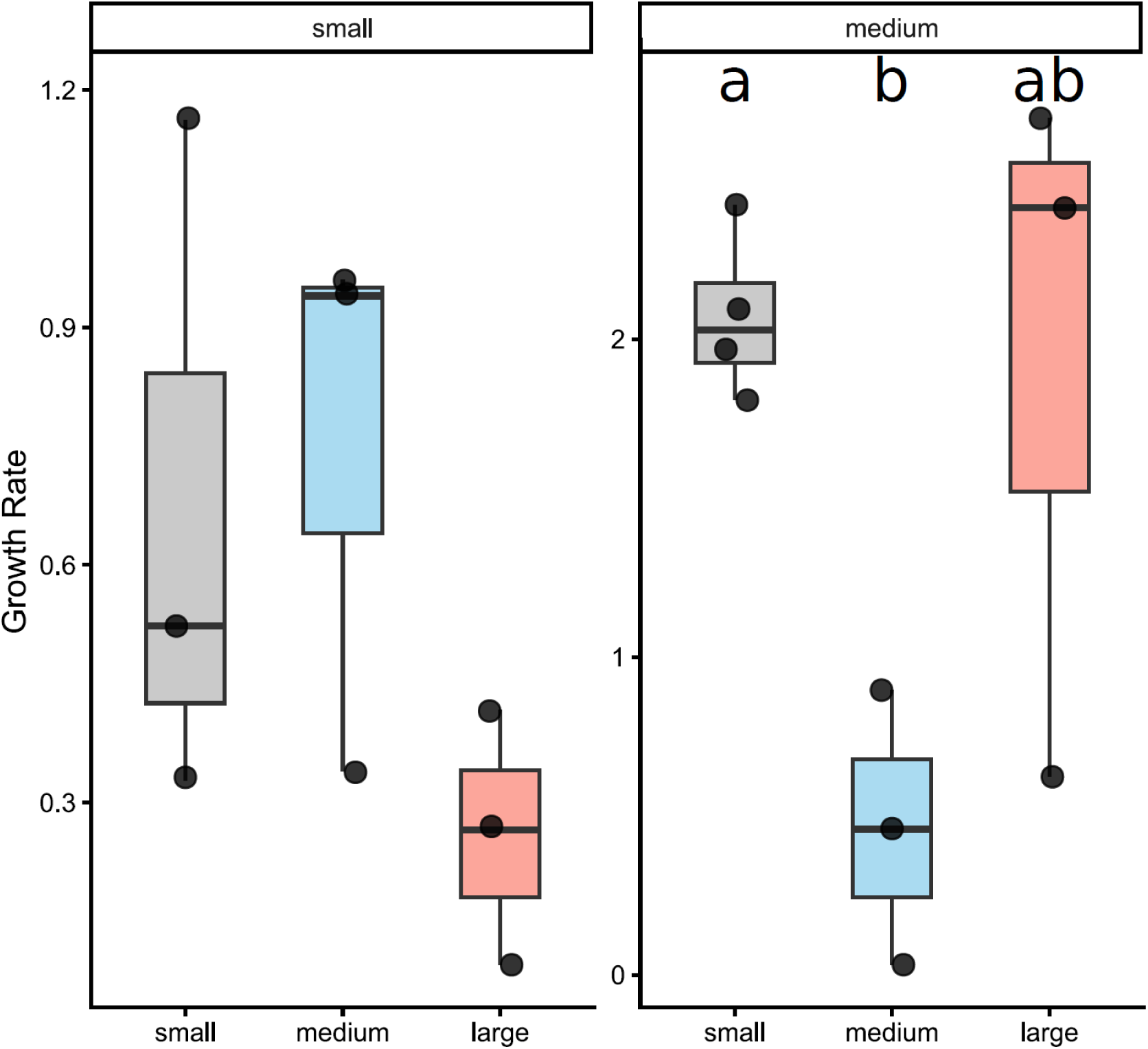
Growth rate of F1 field-derived worms: small and medium-sized worms exposed to differently sized worms over two weeks. As previously, exposure to small worms served as control. Additionally, worms were exposed to medium-sized and large (adult) worms. The boxplots depict the median growth rate (segment length grown/week) with the first and third quartile of the distribution. Whiskers extend to 1.5 × the interquartile range; data beyond that range are defined as outliers and plotted individually. The differences between the difference in growth under different conditions for the medium worms are indicated by letters, defined by Tukey post-hoc tests.

Only in the experiment using medium worms from F1 field-derived culture, we measured a significant effect of the condition (small, medium, large) on the growth rate (Anova, F=5.68, 𝑃 = 0.034; Fig S3). With Tukey post-hoc tests, we measured a significant reduction in the growth rate in medium worms exposed to other medium worms (mean growth rate= 0.46 +- 0.43 segments/week) compared to those exposed to small worms (mean growth rate= 2.07 +- 0.26 segments/week; Tukey post-hoc test, p.adj=0.037; Fig 7). The growth rate of those exposed to large worms (mean growth rate= 1.91 +- 1.12 segments /week) was not significantly different from either of the two other conditions (exposed to small, exposed to medium). This experiment indicates that medium worms (>40 segments) also significantly stunt juvenile growth.

### Chemical Characteristics of Growth Suppression (c)

In order to characterize the chemical cue(s) responsible for the observed growth suppression, we first tested whether they originate from faeces. There was no significant difference in growth rate (Welch t-test, t=-1.189, df=2.815, 𝑃=0.3249; Fig. 8; Fig S4) in worms exposed to adult worm faeces (mean growth rate= 3.53 +-0.72 segments/week) compared to those exposed to clean culture water (control, mean growth rate= 2.98 +-0.34 segments/week).

**Fig. 8.**
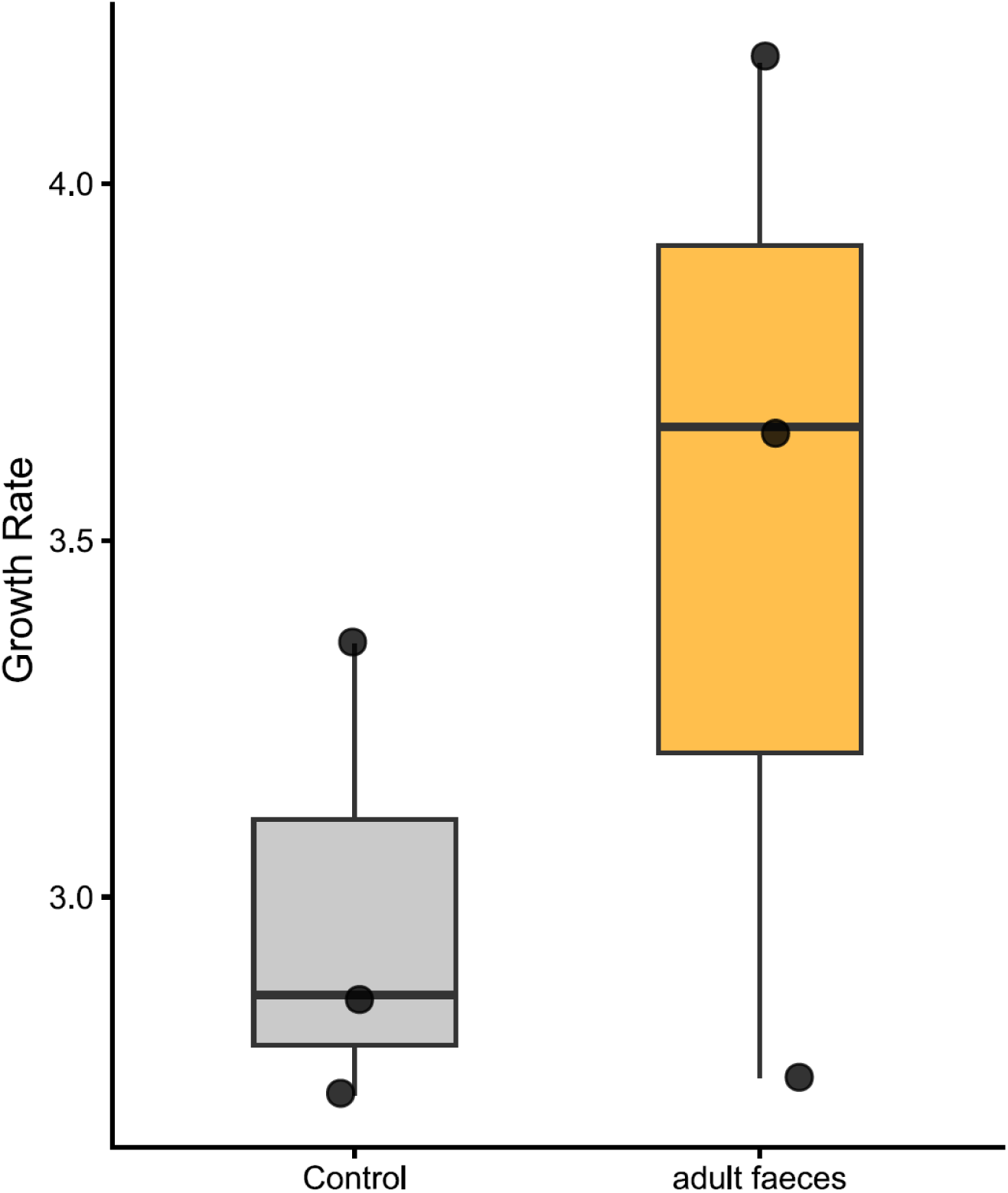
Growth rate of worms exposed to clean culture water (control) compared to exposure to adult worm faeces over the course of 3 weeks. The boxplots depict the median growth rate (segment length grown/week) with the first and third quartile of the distribution. Whiskers extend to 1.5 × the interquartile range; data beyond that range are defined as outliers and plotted individually. There is no significant difference between the treatment groups according to Welch t-test.

Whilst adult worm faeces had no effect on the growth rate of juveniles, we measured a significant difference between the growth rate of worms exposed to water from adult worm tanks (mean growth rate=0.56 +- 0.32 segments/week) and clean culture water (mean growth rate= 4.72 +- 0.36 segments/week; Welch t-test, t=17.9, df=3.21, 𝑃=0.0003; Fig. 9; Fig S5).Hence, our results in separate tank conditions (Fig. 9) coincide with our results from experiments with the tank dividers (Fig. 6), confirming that water-borne chemicals excreted by adult worms have the potential to stunt the growth of juveniles.

**Fig. 9.**
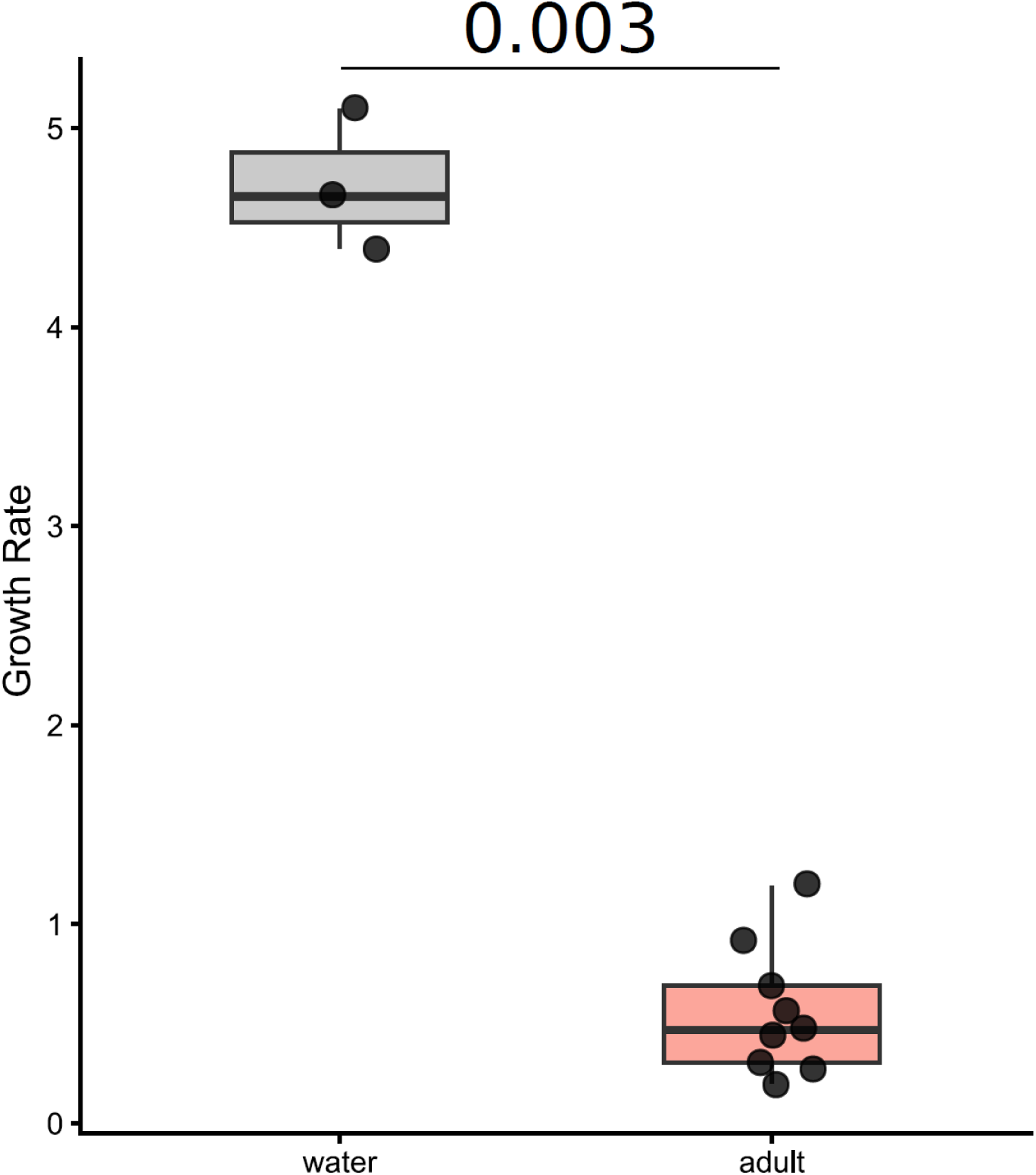
Growth rate of small worms exposed to clean culture water (control) and water from adult worm tanks over 3 weeks. The boxplots depict the median growth rate (segment length grown/week) with the first and third quartile of the distribution. Whiskers extend to 1.5 × the interquartile range; data beyond that range are defined as outliers and plotted individually. The p-value of the Welch t-test comparing the two conditions is written between the two boxes.

To determine some characteristics of the chemical cue, small worms were exposed to water fractionated by size (3 kDa). This allowed us to explore whether the active growth suppressing compound is smaller or larger than 3 kDa. We measured a significant effect of the condition to the growth rate (Anova, F=107.7, 𝑃=0.002; Fig. 10). Indeed, the growth rate of those exposed to <3 kDa was significantly lower being ca. half reduced (mean=1.78 +- 0.11 segment/week) compared to worms exposed to control water (mean=3.65 +- 0.28 segment/week; Tukey, post-hoc test, p.adj=0.003), and that those exposed to >3 kDa (mean=3.93 +- 0.04 segment/week; Tukey, post-hoc test, p.adj=0.002; Fig. 10; Fig S6).

**Fig. 10.**
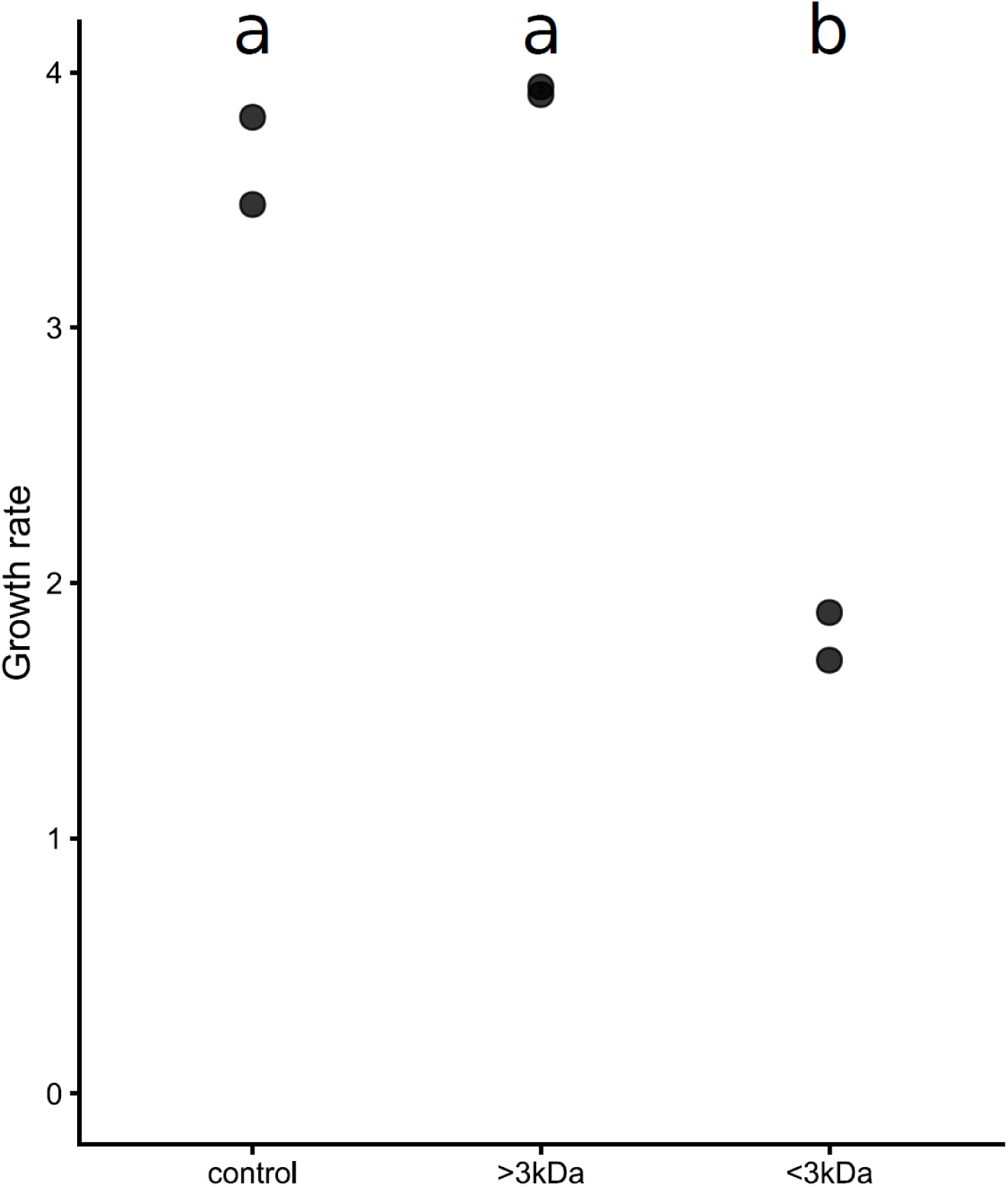
Growth rate of small worms exposed to clean culture water (control) and to >3 kDa and <3 kDa extracts of adult worm water over the course of 2 weeks. The differences in segment growth under different conditions for the small worms are indicated by letters, defined by Tukey post-hoc tests.

## Discussion

Studying growth of *P. dumerilii* worms in the presence of differently sized conspecifics improved our understanding of how hierarchy systems and dominance can be established. Our data suggest that *P. dumerilii* engages in interference competition by chemically suppressing the growth of conspecifics, a non-consumptive mechanism whereby individuals reduce others’ access to resources without direct consumption (Case and Gilpin, 1974). Across experimental conditions, juveniles exposed to larger worms consistently show reduced segment growth compared to controls, demonstrating a clear and repeatable suppression effect.

Although absolute growth rates varied between treatments, the direction of the effect remained consistent. Worms from the established laboratory culture showed strong growth reduction in the presence of larger individuals, whereas F1 field-related worms exhibited overall lower growth, likely due to stress from environmental transition (temperature, salinity, food availability) after only one laboratory generation. Despite this, the same qualitative pattern persisted: larger individuals suppressed growth of smaller and intermediate-sized worms. With F1-field collected individuals, medium worms exposed to medium conspecifics showed significant growth reduction, indicating suppression already occurs above ∼40 segments. Non-significant differences in other comparisons using F1 field-related worms likely reflect reduced growth and limited sample size rather than absence of effect. Experimental separation confirms that growth suppression occurs without direct contact, indicating mediation by diffusible chemical cues and placing the interaction within chemically mediated interference competition.

A key feature of the mechanism is size dependence. Juvenile *P. dumerilii* grow ∼one segment per day (Hauenschild and Fischer, 1969), enabling fine-scale comparisons across size classes. Individuals above approximately 40 segments (medium and large) consistently induce growth suppression in smaller worms, while smaller individuals do not exert comparable effects. This suggests the presence of a size-dependent threshold, beyond which individuals acquire the capacity to influence the growth of others. Notably, this threshold corresponds closely to developmental transitions in *P. dumerilii*, including the onset of gametogenesis and changes in growth dynamics (Fischer et al., 2010; Kuehn et al., 2022). Such alignment supports the hypothesis that the production of the suppressive cue is linked to maturation state. In that case, the suppressive cue may arise as a by-product of maturation-linked physiological changes. Metabolic and hormonal transitions may generate diffusible compounds affecting nearby individuals, as waste products can function as chemical cues in aquatic systems (Brönmark and Hansson, 2012).

Across taxa, similar size-dependent growth inhibition is widely observed. In amphibian larvae (*Rhinella marina*, *Rana temporaria*), gastropods, and crustaceans (*Macrobrachium rosenbergii*, *Homarus* spp.), larger individuals suppress growth of smaller conspecifics via waterborne cues (Wong et al., 2000; Crossland and Shine, 2012; Pearce, 1997; Juarez et al., 1987; Nelson and Hedgecock, 1983). Our findings of chemically-mediated growth suppression in *P. dumerilii* are comparable to growth inhibition in lobsters (*Homarus*), where a rapidly degrading chemical cue has been shown to inhibit growth in a highly size-dependent manner (Nelson and Hedgecock, 1983). Interestingly, the inhibitory compound in lobsters is not species-specific. This raises important questions about the nature of the chemical cues in *P. dumerilii*. Notably, both annelids and crustaceans share the use of the juvenile hormone methylfarnesoate in growth regulation (Borst et al., 1987; Schenk et al., 2016), further supporting potential mechanistic parallels. Future research should aim to identify the active chemical cue in *P. dumerilii* and determine whether its effects are species-specific or broadly active across related or co-occurring taxa.

Chemical characterization here could already show that the cue is water-soluble and stable for several days based on the experiment exposing juveniles to conditioned water from adult tanks (changed every 3 days). This method excludes potential visual effects and direct contact of juveniles and cue donor individuals as well as further build-up of waste or worm produced metabolites. These findings contrast the rapidly decaying compound responsible for the growth suppression in lobsters (Nelson and Hedgecock, 1983). Fractionation indicates an active component <3 kDa, consistent with small peptides or other low-molecular-weight compounds. Assuming an average amino acid weight of 110 Da, this peptide or protein would be smaller than 27 amino acids, although non-peptidic molecules of similar size cannot be excluded. Proteins and peptides are common in chemical communication (Hardege et al., 2004; Rittschof et al., 1990; Wyatt, 2014), and offer a huge variety of chemical signaling functions.

It is possible that the observed effect results from a combination of compounds acting synergistically, rather than a single active molecule. This interpretation is supported by the extensive changes in gene expression and neuropeptide signaling in the brain of nereidids during maturation (e.g. Conzelmann et al., 2013; Schenk et al., 2019). In addition, specific neuropeptides of the corazonin/GnRH superfamily have been shown to influence growth and maturation processes in *P. dumerilii* (Andreatta et al., 2020), supporting the idea that defined molecular pathways are associated with the maturation state. While these studies do not identify a secreted factor mediating inter-individual communication, they indicate that maturation is characterized by distinct molecular signatures that could give rise to diffusible cues. Further work will be required to determine their identity and whether multiple components contribute to the observed effect.

The presence of chemically mediated growth suppression offers an explanation to the differences in size distributions in communal and individual cultures. In communal cultures, worms exhibit broad and often multimodal size distributions, with some individuals growing rapidly while others remain stunted. Larger individuals inhibit the growth of smaller ones, thereby maintaining or amplifying size differences over time. Already after 2 weeks, worms cultured communally were found to have a broader size distribution than individually cultured worms (Fig. 3a vs b). Growth suppression of smaller worms by larger worms in the communal tanks could explain why, under comparable space and food conditions, communally cultured worms grow less uniformly than individually cultured worms.

The potential of early social imprinting might explain why we observed several peaks at the beginning of our size distribution experiments. The worms already show a small difference in their growth rate with some jumpers growing faster (Fig. 3: peaks on the right side of the mean) and others growing slower (Fig. 3: peaks on the left side of the mean). Indeed, after six weeks, both communal and individually reared *P. dumerilii* displayed bell-shaped size distributions with early divergence into fast-growing “jumpers” and stunted “laggards,” independent of initial size (Fig. 3). Although both rearing conditions show evidence for this divergence, under comparable space and food conditions, the effect is markedly stronger under communal conditions where overall population growth is reduced compared to individually cultured worms. This mirrors findings in prawns and amphibians where early social environments shape individual growth trajectories (Karplus and Hulata, 1995; Monaghan, 2008; Morey and Reznick, 2000), but requires experimental proof in *P. dumerilii* in future studies.

The release of growth-suppressing cues by adults in *P. dumerilii* likely functions as a density-dependent regulatory mechanism, curbing juvenile development in crowded populations (Light, 1967; Rosenheim and Schreiber, 2022). In natural environments, this may limit growth under high density and influence spatial distribution, as juveniles may avoid areas with high concentrations of larger conspecifics through active movement or selective settlement, thereby reducing exposure to inhibitory cues. Field and multi-generational culture observations support this idea, as individuals from the same clutch often show strongly divergent growth trajectories, reaching maturity over 3 to 12+ months (Hardege, pers. comm.). In aquatic systems, size-structured competition frequently generates similar outcomes, with larger individuals suppressing or outcompeting smaller ones, and in some taxa even consuming them—e.g., amphibians (Crossland and Shine, 2011; Pfennig, 1990; Pfennig, 1999) and fishes (Jonsson, 2025; Polis, 1981). Claessen et al. (2000) further showed that such asymmetric interactions, including cannibalism, can produce “giant” and “dwarf” phenotypes, highlighting how feedbacks in size-structured systems amplify divergence within cohorts.

Although cannibalism has not been documented in *P. dumerilii*, aggressive interactions are frequently observed, and cannibalism occurs when individuals of different sizes are housed together. Chemical cues from conspecific or released during cannibalistic interactions can be used to avoid conspecifics in cannibalistic species and potentially mediates complex behavioral trade-offs between feeding opportunities and predation risk. For instance, odors from injured blue crabs (*Callinectes sapidus*) deter conspecifics from conflict zones (Moir and Weissburg, 2009), whereas hermit crabs (Paguroidea) increase foraging activity in response to similar cues (Tran, 2014). In damselfly larvae (*Ischnura elegans*), cues released by larger conspecifics signal danger or competition, eliciting reduced activity, greater refuge use, prioritizing survival over foraging (Sysiak et al., 2023). In *Platynereis dumerilii*, chemical suppression may function analogously, allowing dominant individuals to outgrow others without physical elimination. Like cannibalism, this may evolve under high density or limited resources (Rosenheim and Schreiber, 2022). such dominance mechanisms can create bimodal size distributions, as seen here with *P. dumerilii* in the first experiment, in fish (Jonsson, 2025), and in amphibians (Griffiths et al., 1991), where stunted individuals may shift niches or remain subordinate. The persistence of suppressed growth in *P. dumerilii* suggests lasting developmental consequences, potentially akin to plastic “dwarf” phenotypes described in other taxa (Monaghan, 2008).

In natural systems, these mechanisms may also influence ecosystem dynamics. *P. dumerilii*, as well as other Nereididae, play a fundamental role in ecosystem functioning (Lawrence and Soame, 2009). As grazers and preys, they are vital components of food webs in marine and estuarine environments. The role of chemically mediated growth suppression on ecosystem dynamics remains to be discovered. Interestingly, in the field, monthly reproduction is observed. This corresponds to laboratory conditions with growth in batches where large individuals (and their chemical stunt cues) are removed once a month, when ripe heteronereids reproduce and die (see Özpolat et al., 2021, for review), enabling medium sized worms to grow and metamorphose for the next month’s spawning event.

From an applied perspective, these findings are directly relevant to laboratory culturing. In communal tanks, increased mortality may result not only from growth suppression but also from increased disease transmission due to infrequent water changes (to maintain chemical cues), as well as fighting, all of which negatively impact health and segment growth (Evans, 1973). To minimize these effects, cultures could be rapidly expanded by maintaining narrow size distributions and regularly removing large individuals. This would promote uniform growth and increase yield, improving *P. dumerilii’s* utility as a model organism for chronobiology and developmental biology (Kuehn et al., 2019). Our findings reinforce that chemical interactions must be considered alongside food and space in rearing protocols.

As growth suppression is known to occur in other marine species (most notably crustaceans (Karplus and Hulata, 1995; Karplus et al., 1992; Nelson and Hedgecock, 1983)), improving our understanding of the mechanisms underlying growth suppression in *P. dumerilii* could have major implications for aquaculture conditions. Differential growth is indeed a known problem in aquacultures, where high population densities are common. Our study opens up new avenues of research, addressing chemical interactions in population dynamics in species living in large communities. Although dominance and hierarchies have been extensively studied in marine environments (Briffa and Sneddon, 2007; Drews, 1993; Gherardi and Atema, 2005), to date, few studies have focused on the role of odour profiles and cue production in these social interactions.

To the best of our knowledge, this is the first demonstration of chemically mediated growth inhibition in annelid worms, offering new insights into non-reproductive chemical communication in marine invertebrates. This highlights the importance of non-lethal chemical interactions in shaping competition, phenotypic plasticity, and population dynamics (Alford and Wilbur, 1985; Narvaez et al., 2020; Wilbur, 1980). Overall, these findings underscore the need to integrate chemical communication into ecological and developmental frameworks and open new avenues for studying the molecular and evolutionary basis of non-consumptive interactions in marine invertebrates.

## Acknowledgements

The authors would like to thank Margaret Huffee for technical support, Maggy Harley for advice and help with cue purifications, and Prof. Gaspar Jekely for providing the worms. We also acknowledge Dr. Helga Bartels-Hardege for advice and feedback on the manuscript.

## Data and resource availability

All data is deposited in the supplementary information. All collections and experiments were conducted following national and/or institutional guidelines. Approval was granted by the Ethics Committee of the University of Hull (No. UO20).

