## Supplementary Figures for "Adult Marine Annelid *Platynereis dumerilii* Chemically Stunt the Growth of Juveniles"

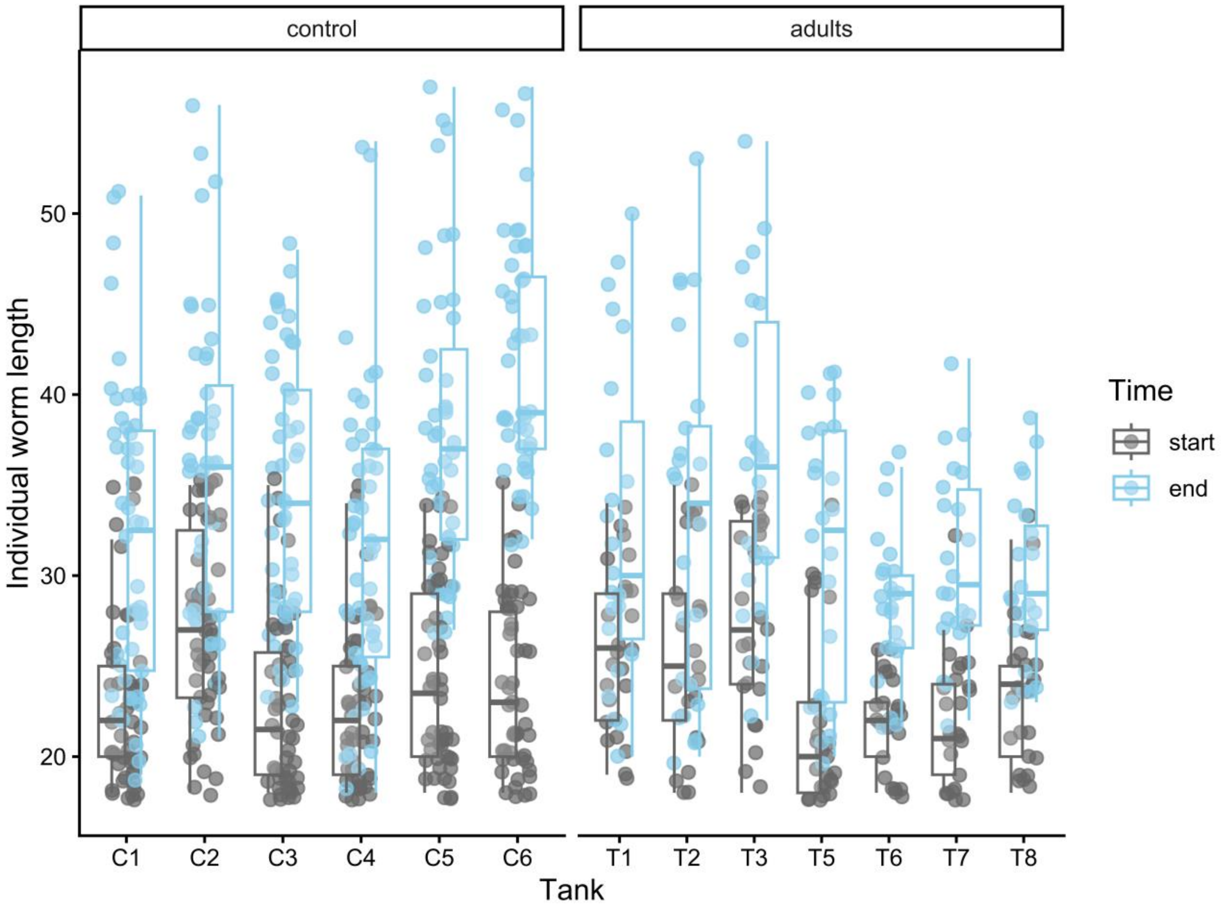


**Fig S1** Individual worm length (number of segments) of small worms from an established laboratory culture exposed to small worms (control) or large worms (adults), measured before exposure (start, grey) and after 3 weeks (end, light blue). Each dot represents an individual worm measured within its tank. The boxplots show the median segment length within each tank, with the first and third quartiles. Whiskers extend to 1.5 times the interquartile range, and data beyond this range are considered outliers and plotted individually.

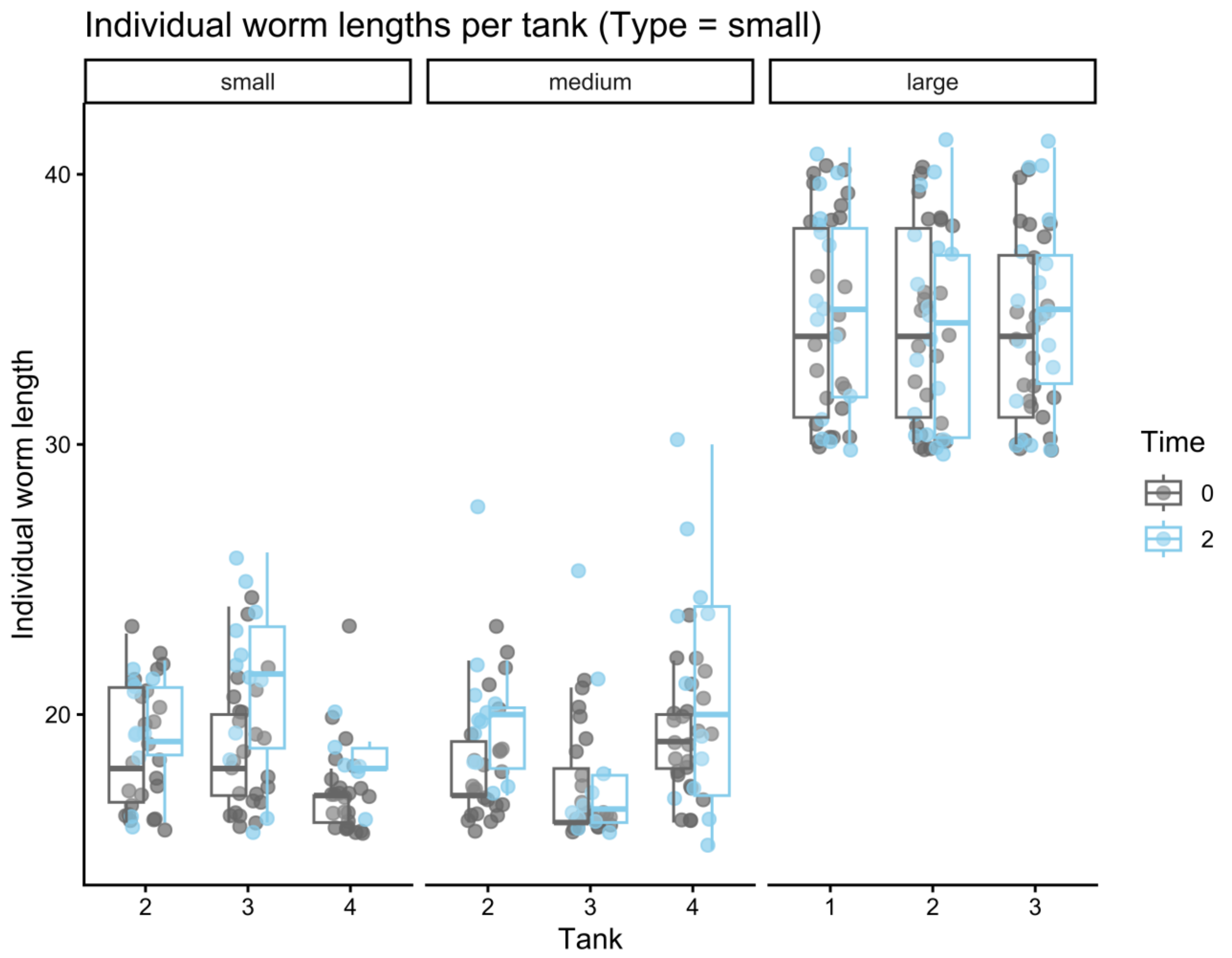


**Fig S2** Individual worm length (number of segments) of small worms collected in the field exposed to small worms (control), medium or large worms (adults), also collected in the field, measured before exposure (0, grey) and after 2 weeks (2, light blue). Each dot represents an individual worm measured within its tank. The boxplots show the median segment length within each tank, with the first and third quartiles. Whiskers extend to 1.5 times the interquartile range, and data beyond this range are considered outliers and plotted individually.

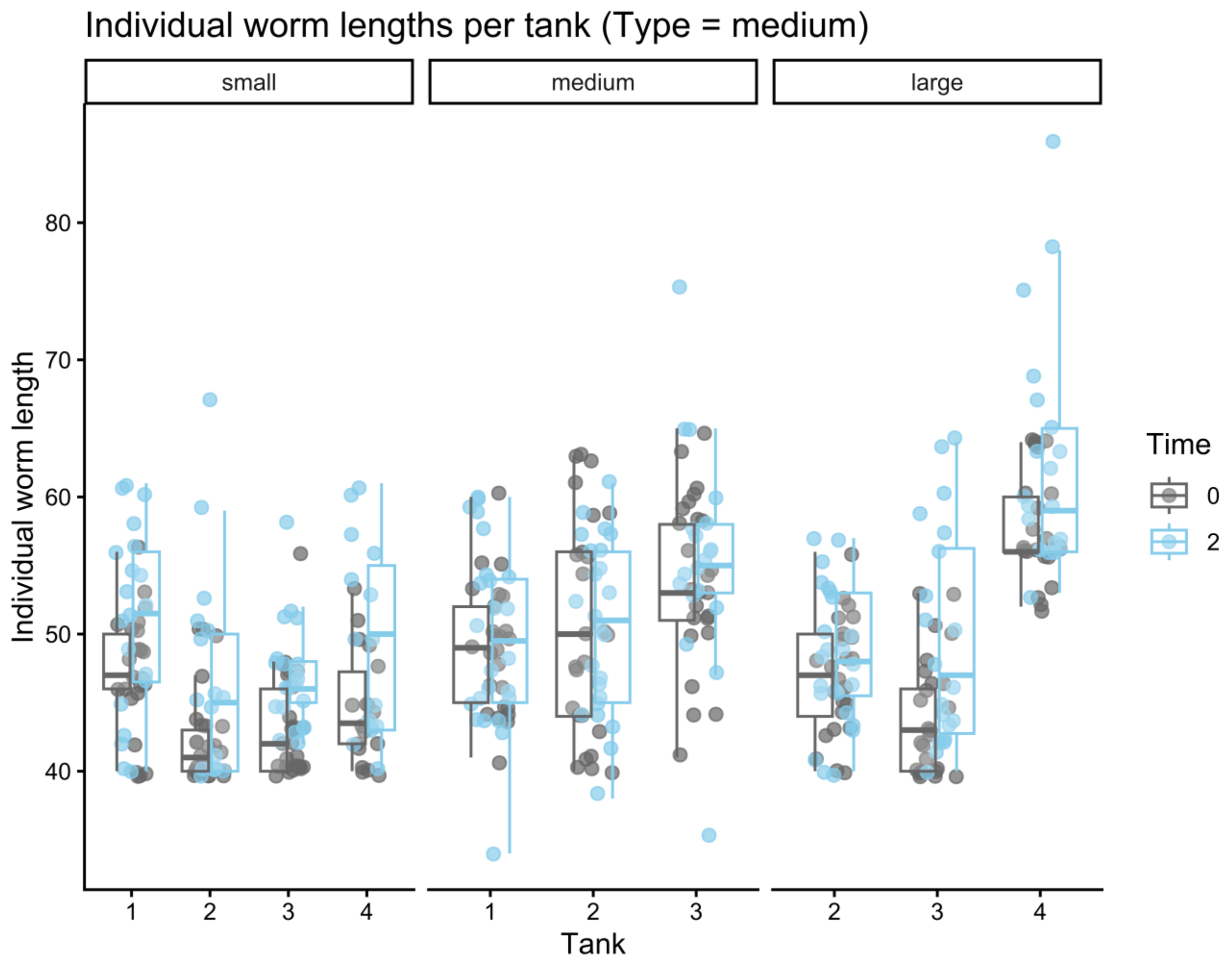


**Fig S3** Individual worm length (number of segments) of medium worms collected in the field exposed to small worms (control), medium or large worms (adults), also collected in the field, measured before exposure (0, grey) and after 2 weeks (2, light blue). Each dot represents an individual worm measured within its tank. The boxplots show the median segment length within each tank, with the first and third quartiles. Whiskers extend to 1.5 times the interquartile range, and data beyond this range are considered outliers and plotted individually.

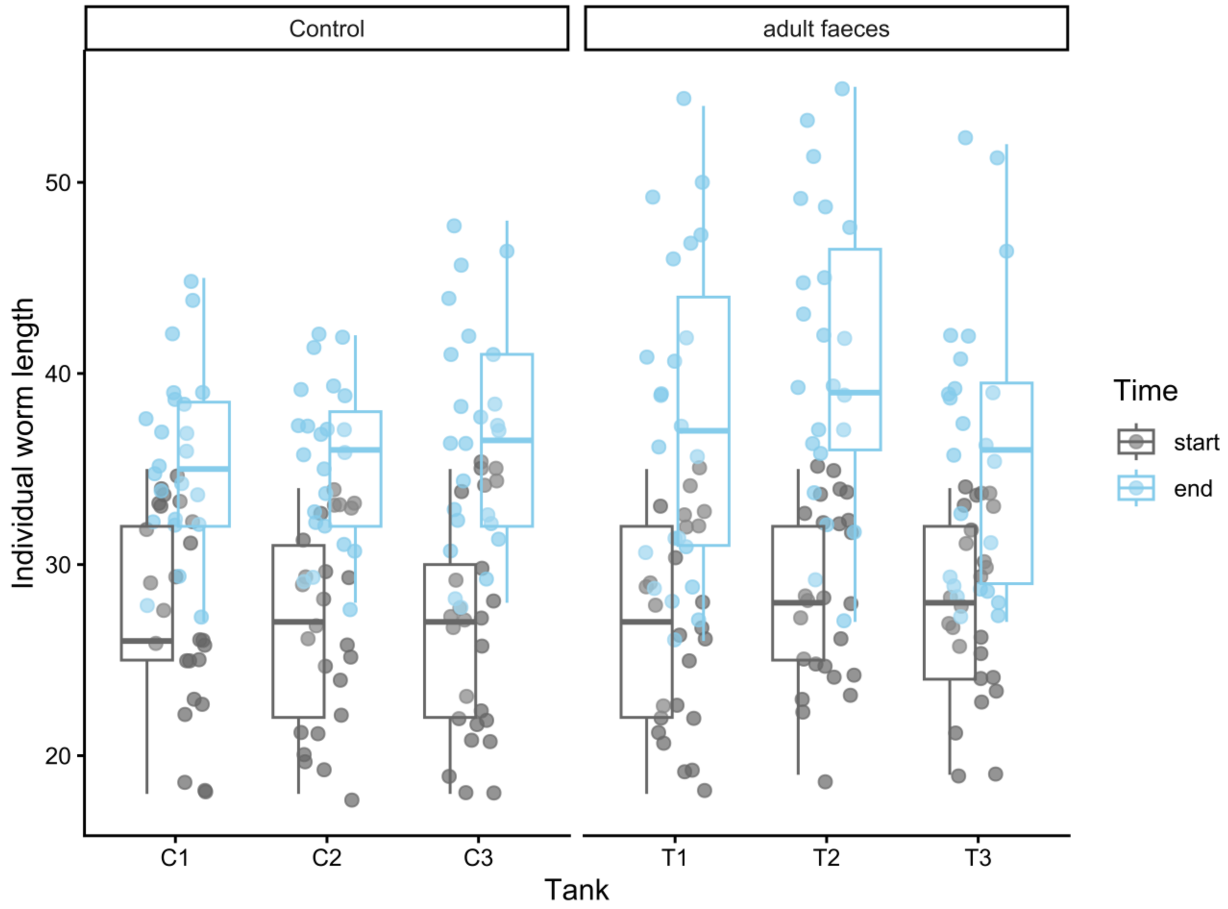


**Fig S4** Individual worm length (number of segments) of small worms from an established laboratory culture exposed to clean culture water (control) or adult faeces, measured before exposure (start, grey) and after 3 weeks (end, light blue). Each dot represents an individual worm measured within its tank. The boxplots show the median segment length within each tank, with the first and third quartiles. Whiskers extend to 1.5 times the interquartile range, and data beyond this range are considered outliers and plotted individually.

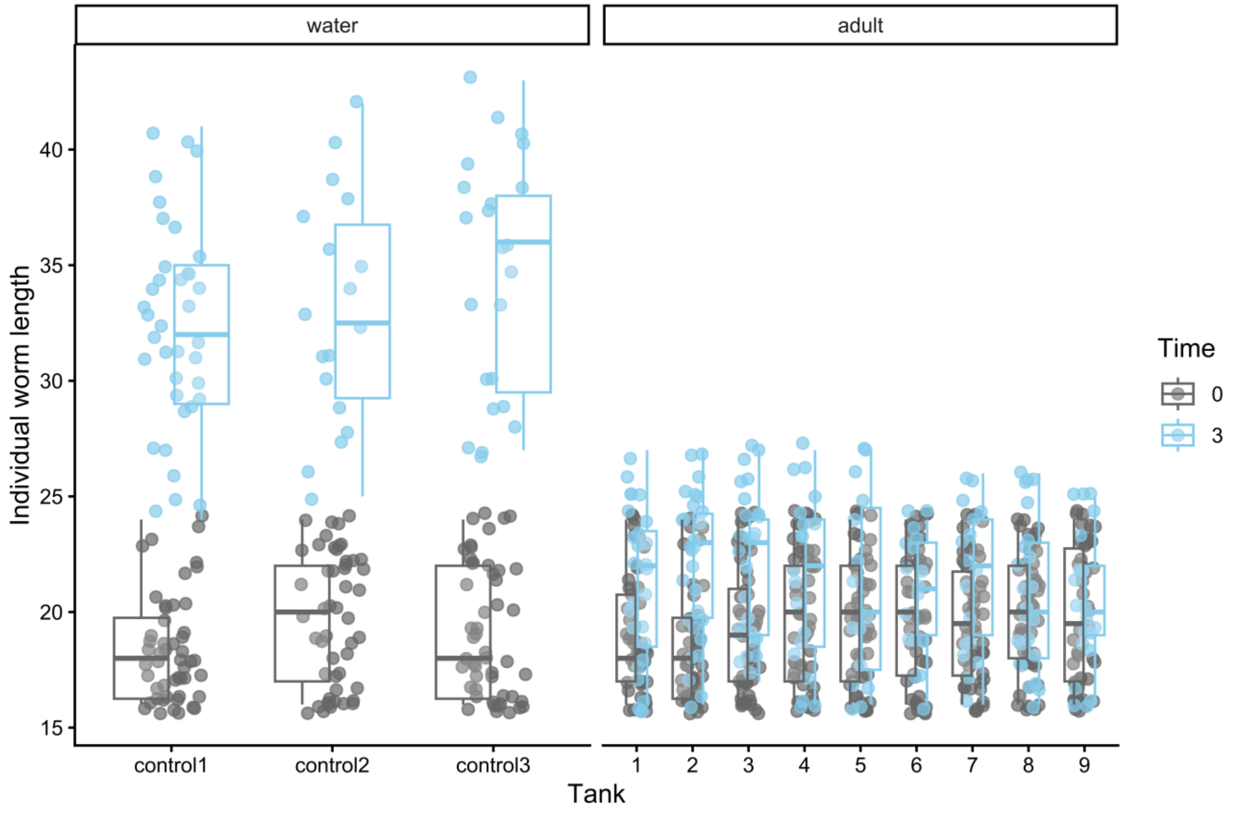


**Fig S5** Individual worm length (number of segments) of small worms from an established laboratory culture exposed to clean culture water (control: water) or adult water (adult), measured before exposure (0, grey) and after 3 weeks (3, light blue). Each dot represents an individual worm measured within its tank. The boxplots show the median segment length within each tank, with the first and third quartiles. Whiskers extend to 1.5 times the interquartile range, and data beyond this range are considered outliers and plotted individually.

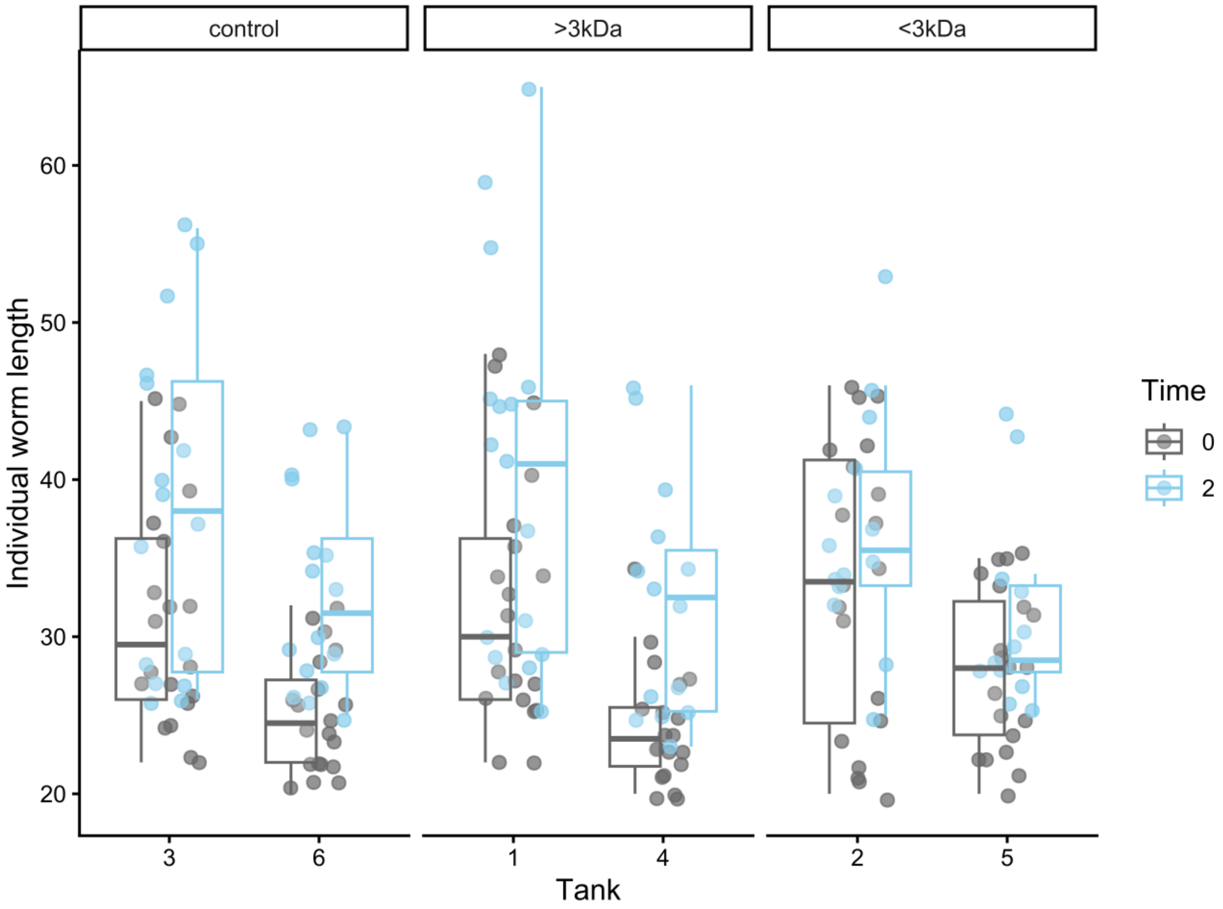


**Fig S6** Individual worm length (number of segments) of small worms from an established laboratory culture exposed to clean culture water (control) and extracts of >3kDa and of <3kDa of adult worm water, measured before exposure (0, grey) and after 2 weeks (2, light blue). Each dot represents an individual worm measured within its tank. The boxplots show the median segment length within each tank, with the first and third quartiles. Whiskers extend to 1.5 times the interquartile range, and data beyond this range are considered outliers and plotted individually.
